# An organ-on-a-chip approach for investigating root-environment interactions in heterogeneous conditions

**DOI:** 10.1101/126987

**Authors:** Claire E. Stanley, Jagriti Shrivastava, Rik Brugman, Dirk van Swaay, Guido Grossmann

## Abstract

Plants adapt their root morphology in response to changing environmental conditions, yet it remains largely unknown to what extent developmental adaptations are based on systemic or cell-autonomous responses. We present the dual-flow-RootChip (dfRootChip), a microfluidic organ-on-a-chip platform for asymmetric perfusion of Arabidopsis roots to investigate root-environment interactions under simulated environmental heterogeneity. Applications range from root guidance, monitoring of physiology and development under asymmetric conditions, tracing molecular uptake and selective drug treatments to local inoculation with microbes. We measured calcium responses in roots treated with biotic and abiotic elicitors and observed elicitor-specific signal propagation across the root from treated to untreated cells. We provide evidence for non-autonomous positive regulation of hair growth across the root upon exposure to unfavourable conditions on the opposite side. Our approach sheds light on lateral coordination of morphological adaptation and facilitates studies on root physiology, signalling and development in heterogeneous environments at the organ level.

## INTRODUCTION

The rhizosphere is a diverse ecosystem and an environment with great structural and compositional complexity. Soil texture and density, as well as its mineral content or oxygen availability, can vary in a way that a single root system may have to adapt to a range of micro-environmental conditions. Due to both biological activity and abiotic conditions, any local micro-environment is permanently subject to changes, rendering rhizosphere conditions highly dynamic. The ability of plants to adapt their root system architecture according to soil conditions is a prime example of developmental plasticity and has attracted increasing attention over the past decades. The emergence of lateral roots and root hairs can be triggered or inhibited in response to environmental stimuli. Soil conditions such as water content, nutrient concentrations or salinity result in characteristic root system architecture through growth modulation of primary and higher order roots as well as the density and length of root hairs (Gruber et al., 2013; Rellán-Álvarez et al., 2016; Williamson et al., 2001). Root hairs are tip-growing protrusions of specialised root epidermal cells that play important roles in soil penetration and absorption of phosphate and other mineral nutrients (Brown et al., 2013; Haling et al., 2013). Root hair development has served as a readout for the root’s response to a lack of mineral nutrients (Bates and Lynch, 1996; Chandrika et al., 2013; Müller and Schmidt, 2004; Song and Liu, 2015). Experiments on root systems exposed to different media conditions showed that both lateral root formation and root hair development are regulated independently of the global metabolic state, but rather responded to local conditions (Bates and Lynch, 1996). The finding that hair development can be suppressed or stimulated on the same root due to variations in local water availability (Bao et al., 2014) highlights the ability of roots to modify their architecture on a cellular level and underlines the need to study developmental adaptation at cellular resolution.

The high sensitivity of root cells towards changing environmental conditions is also illustrated by rapid physiological changes that precede the regulation of root growth and development. Responses to biotic and abiotic stimulation commonly involve calcium signalling (Dodd et al., 2010). The visualisation of cytosolic calcium transients using luminescent or fluorescent sensors has revealed stimulus-specific calcium signatures and long-distance communication in roots (Behera et al., 2015; Choi et al., 2012; 2014; Keinath et al., 2015; Kiegle et al., 2000; Knight et al., 1997; Krebs et al., 2012; Martí et al., 2013; Xiong et al., 2014). Many more studies have highlighted the importance of numerous environmental factors in root development, but how locally perceived signals are communicated to neighbouring cells, how roots differentiate and prioritise between diverse environmental signals and how cell signalling orchestrates root development in complex natural environments is still largely unknown.

A technical challenge in the lab has been the simulation of environmental complexity to reflect the soil conditions that roots are exposed to, whilst at the same time being able to control and quantify this environmental complexity. For high-resolution studies on root signalling and development, plants are commonly grown on synthetic gelled or hydroponic media. This reductionist approach has, however, limitations as environmental diversity may be a critical factor for root development. Indeed, it has been shown that numerous physiological and developmental processes are significantly different between soil-grown and media-grown plants (Downie et al., 2014; Rellán-Álvarez et al., 2015). Hence, there is a lack of knowledge regarding how environmental diversity influences root development. New approaches and technologies are needed that enable root growth in defined, yet increasingly complex, environments with the possibility to locally apply a stimulus to selected cells in a single root.

Organ-on-a-chip technology, i.e. the cultivation of tissues in microfluidic devices, has revolutionised experimental access to animal and plant organs by enabling researchers to precisely control the specimen’s micro-environment and image biological processes within tissues at high spatiotemporal resolution (Bhatia and Ingber, 2014; Sanati Nezhad, 2014; Zheng et al., 2016). This emerging field therefore offers the potential to simulate the natural diversity found in soil. Indeed, the significance of “soil-on-a-chip” microfluidic technologies for investigating complex relationships between soil-dwelling organisms and their environment has been recently highlighted (Stanley et al., 2015).

Herein, we present a microfluidic device, termed the dual-flow-RootChip (dfRootChip), that allows cultivation of Arabidopsis roots in asymmetric micro-environments. We guided root growth through an array of micro-pillars that centred the root and allowed us to generate different conditions on either side of the root using laminar flow. We demonstrate the versatility of the approach by following the uptake of fluorescent molecules, by locally trapping plant-pathogenic bacteria at the root surface and by imaging calcium signals upon selective application of abiotic and biotic stresses. In addition, we studied the role of local environmental conditions on the development of root hairs and found that root hair repression and stimulation can be regulated both cell-autonomously and in a coordinated manner across the root. The dfRootChip therefore provides a means for incorporating environmental complexity into experimental design for studies on plant roots.

## RESULTS

### Design and operation of the dual-flow-RootChip (dfRootChip)

The dual-flow-RootChip (dfRootChip) consists of micron-sized root observation chambers that feature a root guidance array and a perfusion system, mounted on optical glass (**Figure 1A,B**). Each observation channel has a height, width and length of 110 μm, 500 μm and 12 mm respectively. Arabidopsis seedlings are grown through medium-filled plastic pipette tips that are inserted into the root inlet of the observation channel (**Supplementary Figure S1B**). Once the primary root has entered the observation channel via a guidance channel (approx. 7 days after germination, **Supplementary Figure S1**), the length of the chamber permits continuous observation of root growth for 2-3 days, depending on the growth rate of the root. For seedlings (Col-0) grown in the dfRootChip we observed growth rates of 1.7 ±0.4 μm/min (n=15), which falls within a similar range to growth rates observed on gel media (Yazdanbakhsh et al., 2011). A reservoir, connected to the root inlet via a small channel, acts as a means to provide additional medium to each root during initial incubation in growth chambers (**Supplementary Figure S1**). The microchannels were engineered to have a target height of 110 μm as this is the approximate diameter of an Arabidopsis root. Hence, the growing root is confined in the z-direction, which aids imaging of the roots. The compact design of the dfRootChip enables Arabidopsis seedlings to be mounted and several experiments to be performed in parallel. We exemplify this by combining five chambers in parallel on a single glass slide (**Figure 1A**).

**Figure 1.**
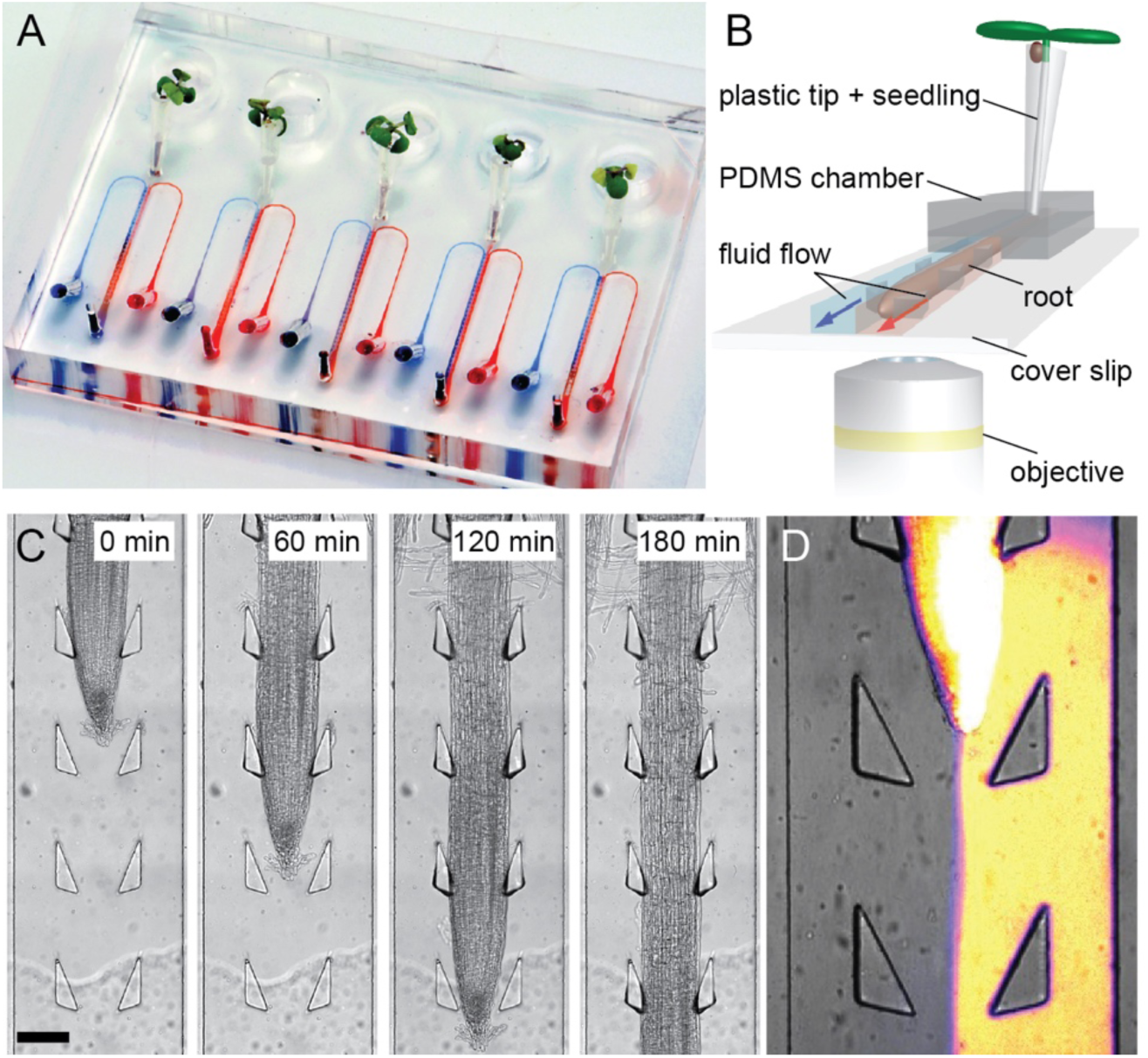
The dual-flow-RootChip (dfRootChip). (**A**) Photograph illustrating the dfRootChip mounted with five Arabidopsis seedlings. Two inlet channels serve each chamber and are coloured red and blue. (**B**) 3D schematic of the device illustrating the direction of flow within the two microchannels. The blue and red colours indicate the ability to deliver two different reagents simultaneously to either side of a growing root. (**C**) Time-series illustrating growth and guidance of an Arabidopsis root through the array of flexible pillars (see also **Supplementary Movie 1**). Scale bar, 100 μm. (**D**) Generation of an asymmetric micro-environment using a fluorescein-containing solution for visualisation purposes. At these length scales laminar flow dominates, demonstrated by the presence of a boundary between the two co-flowing liquids.

One of the major components of the dfRootChip is the root guidance array, which consists of sets of triangular-shaped micropillars distributed in a regular arrangement throughout the root observation channel (**Figure 1C**, **Supplementary Figure S1**). This design specifically enables the direction of root growth to be controlled by centring the root tip in the root observation channel (**Figure 1C**). An inter-pillar distance of 355 μm (along the observation channel) was chosen as this corresponds to a size that is slightly smaller than the distance between elongation zone (EZ) and Arabidopsis root apex. This distance prevents the touch-sensitive columella cells from being stimulated by the pillars and thereby avoids excessive bending of the flexible young shaft, hence avoiding divergence of the root from the desired path. A triangular shape was found to be an appropriate pillar design for aiding the root tip through the pillar array; the shallow angle of the triangular pillar guides the growing root tip towards the centre of the observation channel (**Figure 1C**, **Supplementary Movie 1**). As the microfluidic devices are comprised of the elastomeric polymer poly(dimethylsiloxane) (PDMS), the tips of the micropillars are flexible and deform when the growing root passes them. Hence, the micropillar array does not only act as a guidance mechanism, but also more accurately simulates the presence of loosely packed soil particles.

Two inlet channels and one outlet serve each observation chamber (**Figure 1A, Supplementary Figure S1**). Laminar flow is dominant at these length scales, with a boundary between two co-flowing liquids being evident beyond the root tip (**Figure 1D, Supplementary Movie 2**). Consequently, the dfRootChip design allows a growing root to be subjected to two different flow conditions simultaneously by utilising laminar flow to generate separate micro-environments, enabling either symmetric or asymmetric perfusion of the root. The guidance channel, which connects the “root inlet” to the observation channel, prevents fluid escaping into the reservoirs and therefore disruption of the channel flow.

### Tracing uptake of small molecules into roots

To test to what extent stable asymmetric availability of small molecules is reflected by gradients inside the root we subjected roots grown inside the dfRootChip to 1 μg/ml fluorescein on one side and traced the uptake of the fluorescent dye over time (**Figure 2A,B**). Within 30 seconds, we observed a fluorescence signal first in root hairs, followed by a signal increase in cortical tissues on the side of treatment. The signal intensities inside cells substantially exceeded the signal of the treatment solution, which may indicate dye accumulation inside cells, but can also be explained by pH-dependent quenching of fluorescein fluorescence in the plant growth medium used (1/2x Hoagland’s medium, pH 5.7) (1/2x HM). During the course of dye application (8 min), the signal inside the root did not spread evenly across the organ but instead remained largely retained to the treated side, which indicates that it is possible to generate root-internal gradients and trace the uptake of small molecules into the root. The degree of retention appears, however, dependent on the root zone. Consistent with a maturing endodermis that becomes sealed by the Casparian strip, the signal spread only slowly beyond the treated cortical tissues in the differentiation zone (DZ) and spread more rapidly in the EZ and in zones towards the root tip (**Figure 2A**, last panel).

**Figure 2.**
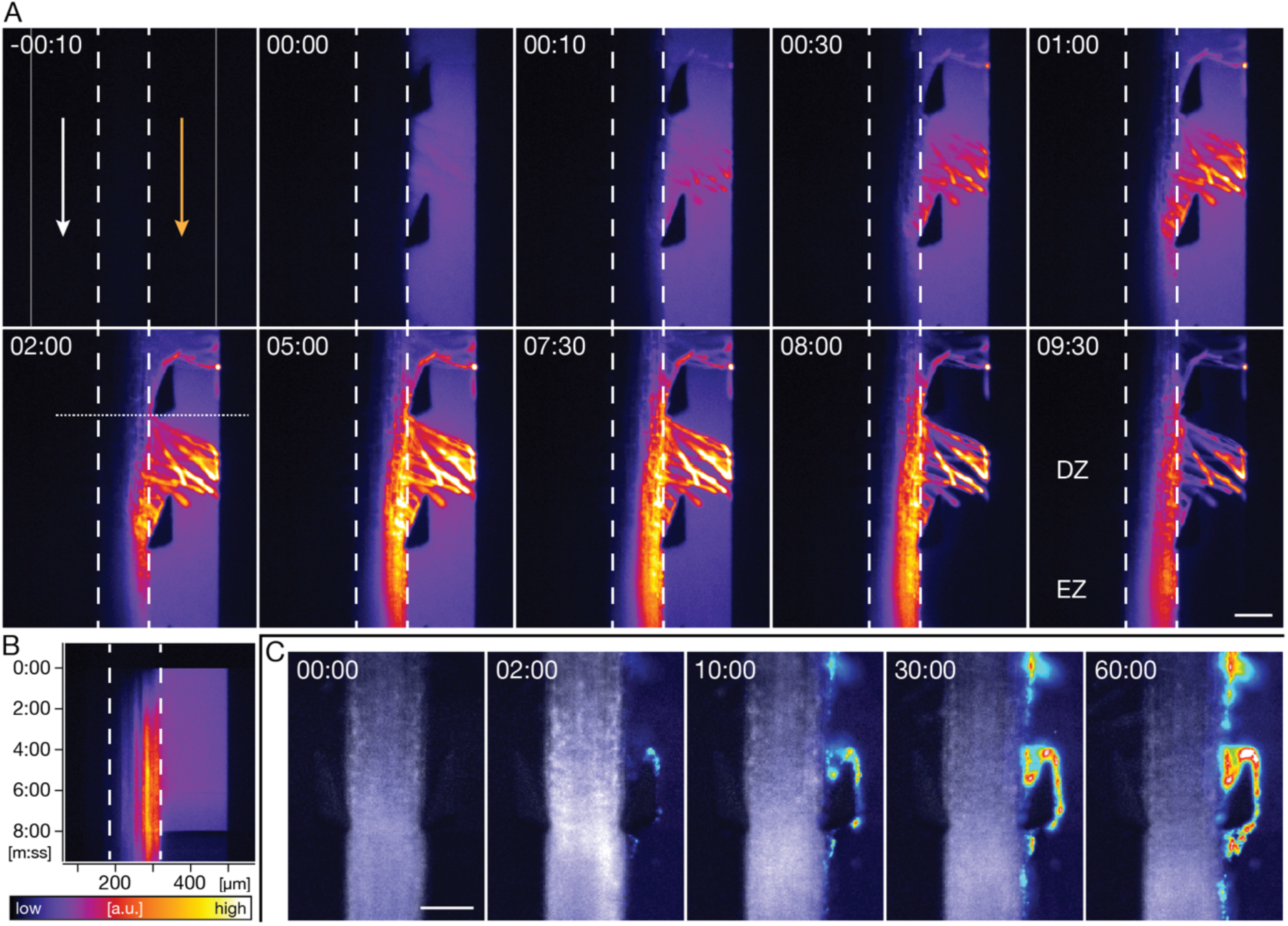
Establishing asymmetric root micro-environments in the dfRootChip. (**A**) Uptake of fluorescein upon asymmetric perfusion of an Arabidopsis root over time (time stamp format, mm:ss). The dashed lines outline the root boundaries. Arrows depict the flow direction of 1/2x HM; the orange arrow indicates the channel where a pulsed treatment using a fluorescein-containing solution (1 μg/ml fluorescein in 1/2x HM) was performed on one side of the root between time points 0 and 08:00. (**B**) Kymograph generated along the dotted line displayed in the image above (panel 02:00 in (**A**)). (**C**) Time series of the association of GFP-expressing *Pseudomonas fluorescens* using the triangular pillars as traps. Scale bars, 100 μm.

### The dfRootChip facilitates investigation of root-microbe interactions

Besides the patchy availability of solutes in natural soil, roots also interact with numerous beneficial or pathogenic microbes, which locally associate with the root. As the nature of this colonisation is for many cases still obscure, there is a high demand for tools that facilitate the visualisation of this process *in vivo* over time. We tested whether the pillar array can be utilised as traps for microbes by inoculating the root with a live culture of the plant pathogen *Pseudomonas fluorescens* expressing green fluorescent protein (GFP) (Haney et al., 2015). Within 10 minutes, we observed a steady accumulation of bacterial cells at the pillar structures, in particular at the pillar tips and in the niche between pillar and root (**Figure 2C**). This shows that the pillar array can indeed be advantageous, when a local increase in microbe density is desired to study the colonisation or infection of roots.

### Selective stimulation with calcium elicitors

The observation that it is possible to generate asymmetric root micro-environments, which result in solute gradients inside the root raises the question to what extent such endogenous gradients are perceived by the root and reflected by intra- and intercellular signalling. Cytosolic calcium elevations are among the first signals that are evoked upon environmental stress and have been used as a readout for cell-to-cell communication in plants (Choi et al., 2014; Dodd et al., 2010). To probe cytosolic calcium elevations ([Ca^2+^]_cyt_) in roots we used Arabidopsis lines expressing the intensiometric calcium indicator R-GECO (Keinath et al., 2015; Zhao et al., 2011). We recorded the [Ca^2+^]_cyt_ response upon selective stimulation of the primary root by performing a pulsed treatment with the elicitor of interest on one side (**Figure 3**). R-GECO-expressing roots grown in the device were exposed to controlled asymmetric perfusion with 1 μΜ flagellin (flg22), a peptide widely used as biotic elicitor that triggers plant immune responses (Felix et al., 1999). The [Ca^2+^]_cyt_ response upon symmetric treatment with flg22 was reported to start in the epidermis and travel inward to the vasculature from where it would travel rootwards and shootwards in the root (Keinath et al., 2015). Upon applying asymmetric treatments we found that [Ca^2+^]_cyt_ only rose in the epidermis on the treated side, after which it propagated into deeper tissues with a velocity of 0.62 ±0.19 μm s^−1^ (n=5) (**Figure 3A, Supplementary Movie 3**). The signal decreased substantially before reaching the untreated side of the root, but, instead, propagated shootwards when reaching the vasculature (**Figure 3B**).

**Figure 3.**
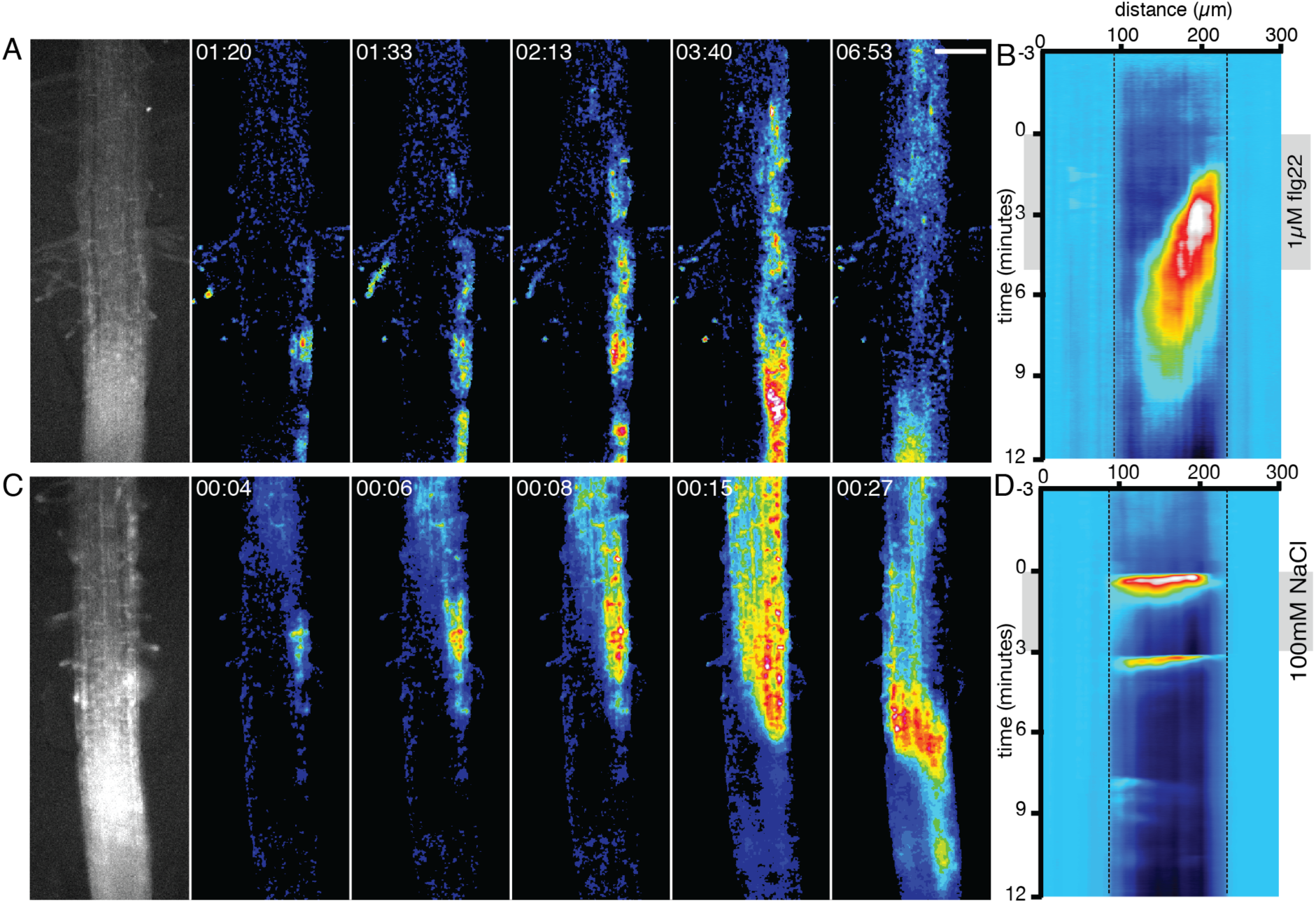
Selective stimulation with calcium elicitors. Calcium dependent signal changes upon asymmetric stimulation with 1 μΜ flg22 (**A, B**) and 100 mM NaCl (**C, D**) in 1/2x HM. Elicitors were applied on the right side using a pulsed treatment, while the left side was perfused continuously with 1/2x HM. (**A**) and (**C**) depict time series of normalised R-GECO fluorescence images (ΔF/F). (**B**) and (**D**) depict kymographs generated from the left to the right of the entire image sequences represented in (**A**) and (**C**) respectively averaging the intensity over the entire height of the image. Scale bar, 100 μm.

To assess the [Ca^2+^]_cyt_ response upon asymmetric exposure to abiotic stress conditions we exposed one side of the root to 100 mM NaCl to mimic salt stress. NaCl treatment is known to generate long-distance [Ca^2+^]_cyt_ signals propagating through the entire root system at velocities of approximately 400 μm s^−1^ when locally applied to the tip of a lateral root (Choi et al., 2014). Under our asymmetric treatments we observed that the [Ca^2+^]_cyt_ signals started in the epidermis on the treated side and propagated across the root to the epidermis on the untreated side at a velocity of 14.1 ±3.61 μm s^−1^ (n=6) (**Figure 3C,D, Supplementary Movie 4**).

Our results demonstrate that the dfRootChip enables selective treatment and recording of [Ca^2+^]_cyt_ elevations in treated and untreated cells simultaneously. Based on the differences in directionality and velocity of calcium signal propagation, we further conclude there must be different mechanisms at play when biotic or abiotic stresses are communicated among root tissues.

### Root hair development in asymmetric growth environments

Since our data support the notion that Arabidopsis roots are able to perceive and process asymmetric stimuli, the question arose as to whether developmental responses and their potential coordination could also be detected upon exposure to asymmetric growth conditions in the dfRootChip. To test whether stimulation of root hair growth is coordinated across the root or regulated cell-autonomously, we compared the effects of asymmetric and symmetric treatments on root hair length in the dfRootChip. One of the best studied conditions leading to altered root hair length is phosphate deficiency (Ma et al., 2001). Numerous studies with seedlings growing on solid media have shown that phosphate deficiency results in longer root hairs, reduced primary root growth and, due to decreased cell elongation, increased root hair density (Bates and Lynch, 1996; Chandrika et al., 2013; Ma et al., 2001; Song et al., 2016). We therefore expected stimulated root hair growth on the phosphate-deficient side. Surprisingly, when we applied different concentrations of phosphate on the two sides of the root, we observed that root hair growth was inhibited and only short bulges (27.8 ±15.5 μm) were formed in trichoblasts on the phosphate-deficient (0.01 mM) side, while normal hair growth was observed on the side supplied with 2.5 mM phosphate (222.0 ±71.3 μm) (**Figure 4B,D**). Under symmetric phosphate conditions, we observed that hair growth was also inhibited under phosphate-deficient conditions (20.6 ±16.0 μm) as compared to symmetric rich conditions (167.9 ±79.8 μm) **(Figure 4A,C,D)**. We obtained similar hair growth phenotypes in experiments using hydroponic growth conditions in medium-filled containers (**Supplementary Figure S2**), which excludes that the unexpected finding was caused by growth in the microfluidic device. Growth tests on vertical plates containing the same medium supplemented with agar revealed, however, stimulated hair development under phosphate-deficient conditions (**Supplementary Figure S2**), which is consistent with published data (Ma et al., 2001).

**Figure 4.**
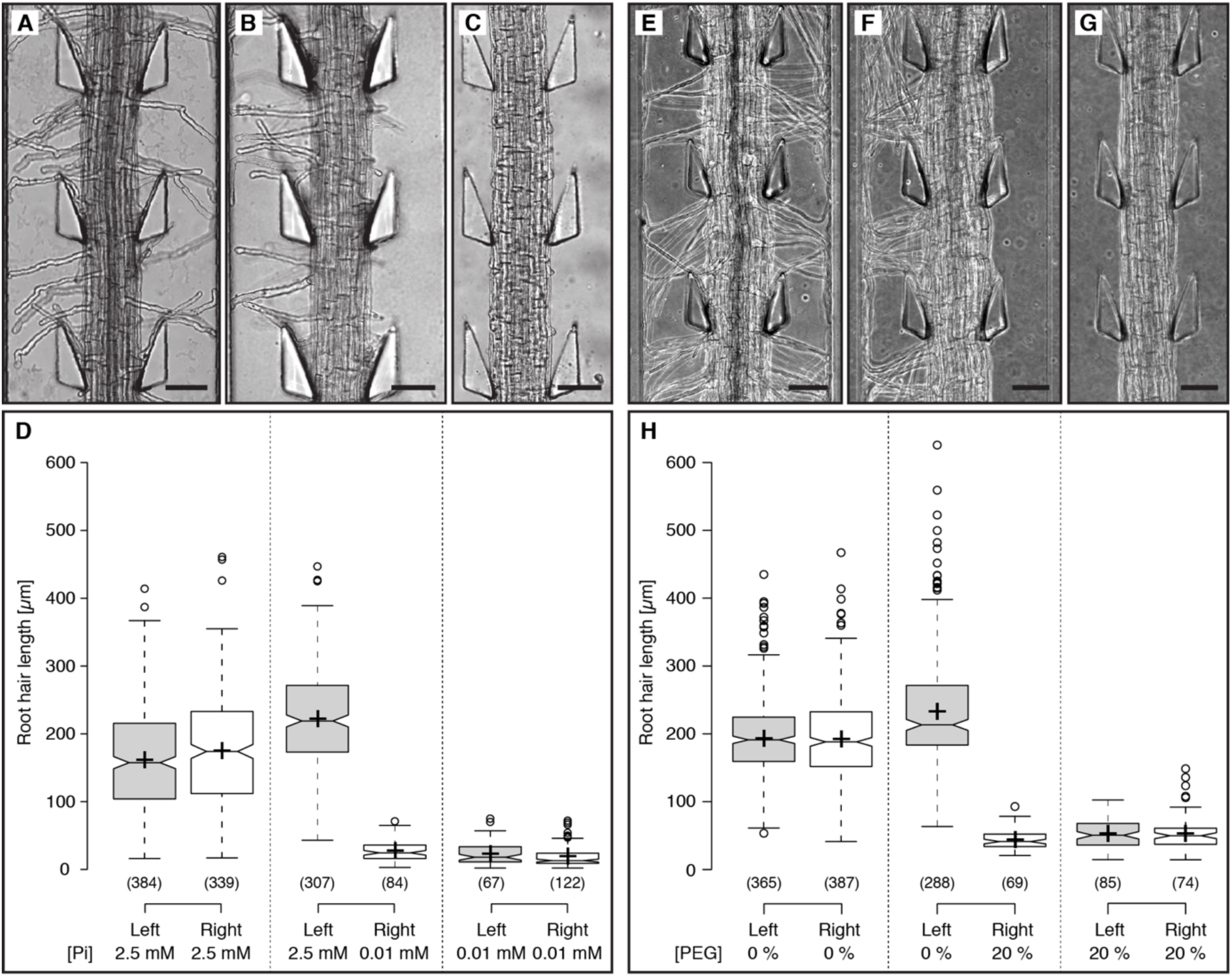
Root hair growth under asymmetric phosphate availability and water stress conditions. (**A-C**) Primary roots of Arabidopsis Col-0 were grown under (**A**) symmetric high phosphate availability (2.5 mM KH_2_PO_4_), (**B**) asymmetric phosphate availability with high phosphate availability on the left side and low phosphate availability (0.01 mM KH_2_PO_4_) on the right side and (**C**) symmetric low phosphate availability. (**D**) Quantification of root hair length for the conditions given in (**A-C**). Centre lines depict the median values, crosses represent sample means; box limits indicate the 25th and 75th percentiles; whiskers extend 1.5 times the interquartile range from the 25th and 75th percentiles, outliers are represented by dots. Sample points (n) from at least 3 independent experiments are given in parentheses below each box. Note that the median values of root hair length under phosphate deficiency are overestimated due to the frequent complete suppression of root hairs or bulges. Only cells with characteristic hair or bulge morphology were included in the analysis. (**E-G**) Roots grown under (**E**) symmetric 1/2x HM, (**F**) asymmetric treatment with polyethylene glycol (PEG) solution (20% (w/v) PEG 8000 dissolved in 1/2x HM), and (**G**) symmetric treatment with PEG solution. (**H**) Quantification of root hair length for the conditions represented in (**E-G**). Box plot annotations are analogous to the graph in (**D**). The overestimation of root hair length under 20% PEG applies for the same reason as in (**D**). Scale bars in (**A-C**) and (**E-G**) represent 250 μm.

When we compared hair length of trichoblasts exposed to 2.5 mM KH_2_PO_4_ under symmetric and asymmetric conditions we noticed an increase in hair length by 32% (*p*=10^−23^) when the opposite side was subjected to phosphate-deficient conditions (0.01 mM KH_2_PO_4_) (**Figure 4D**). This indicates that hair length in cells exposed to rich phosphate conditions can be influenced by the overall phosphate availability.

To test whether the observed asymmetric response in root hair regulation was specific to phosphate availability or a more general phenomenon, we exposed roots to water stress, which has a known inhibitory effect on root hair length (Schnall and Quatrano, 1992). One side of a growing root was subjected continuously to a polyethylene glycol (PEG) solution containing 20% (w/v) PEG 8000 dissolved in 1/2x HM to investigate if root hair growth is influenced upon exerting a low water potential (Ψw), i.e. simulating drought conditions (**Figure 4F**). 1/2x HM containing no PEG was introduced continuously on the other side of the growing root. Typical values for a “low Ψw stress” and “unstressed” conditions have been defined by Verslues *et al*. as a plant cell being exposed to external water potentials equal to -1.0 MPa and -0.2 MPa respectively (Verslues et al., 2006). A 20% (w/v) PEG 8000 solution was chosen as this equates to a water potential of ca. -0.5 MPa (Michel, 1983), thus mimicking low Ψw stress conditions. After symmetric treatment with PEG for 16 hours, the average root hair length was found to be 53.3 ±22.9 μm (**Figure 4G,H**), compared to 193.2 ±62.1 μm under symmetric 1/2x HM (**Figure 4E,H**). We observed that under asymmetric conditions, root hair growth of the side in contact with the PEG solution was arrested (**Figure 4F,H**) (43.8 ±13.5 μm), while root hairs continued to grow on the side containing 1/2x HM only (233.3 ±82.0 μm). Under asymmetric conditions, we further detected a stimulation of hair growth on the 1/2x HM side by 21% (*p*=10^−17^), when compared to symmetric 1/2x HM conditions (**Figure 4H**). This stimulation was in a similar range as observed for asymmetric phosphate availability. Taken together, our experiments on roots under asymmetric phosphate and water availability demonstrate that root hair growth inhibition can be controlled largely cell-autonomously. On the other hand, local deficiencies can stimulate hair length in distal cells, which points to systemic signals or intercellular communication. The mechanisms in charge of these local and distal responses remain, however, to be unveiled.

## DISCUSSION

### The dfRootChip enables high-resolution studies of root interactions with complex micro-environments

Studies on root-environment interactions commonly involve experimental conditions with changes in single parameters to reveal plant responses regarding signalling, metabolism or development. In their natural habitat, plants are, however, exposed to a complex interplay of environmental conditions. Consequently, mechanisms of prioritisation must exist to maximise overall fitness. This includes, for example, the finding that plant immunity responses are suppressed under insufficient light conditions in favour of growth stimulation (Lozano-Durán et al., 2013), or that adaptation of root system architecture to phosphate deficiency depends on the availability of nitrate (Medici et al., 2015). A key to a better understanding of how plants develop in complex environments will be to identify whether responses are systemic or cell-autonomous and to reveal intercellular communication of environmental signals within an organ. It has been recognised that a better understanding of plant developmental plasticity depends on our ability to design experiments that approximate the complexity and dynamics of natural environmental conditions (Rellán-Álvarez et al., 2016).

Here we present an experimental tool that allows the simultaneous stimulation of a single plant organ with spatially confined conditions. The dfRootChip represents an adoption of organ-on-a-chip technology for plants and takes advantage of guided root growth and laminar flow inside the observation channel to create separate environments on both sides of the root. We also show that the triangular pillar design cannot only efficiently guide root growth but is also suitable as traps for bacteria or fungi in host-microbe interaction studies.

Over the past years, microfluidic devices have been developed for plant science to facilitate microscopic access to Arabidopsis roots (Busch et al., 2012; Grossmann et al., 2011; Jiang et al., 2014; Meier et al., 2010; Parashar and Pandey, 2011), pollen tubes (Horade et al., 2013; Nezhad et al., 2013), or moss (Bascom et al., 2016). RootChips have been particularly useful for measuring cellular concentration changes of small molecules using genetically encoded nanosensors for nutrients (Grossmann et al., 2011; Lanquar et al., 2014), phytohormones (Jones et al., 2014) or the second messenger calcium (Denninger et al., 2014; Keinath et al., 2015). While test solutions or changes in the root micro-environment could, so far, only be applied to the root as a whole, the dfRootChip now provides the ability to apply treatments to one side of the root and also study responses in cells that are not in immediate contact with the test solution. In addition, combinations of two treatments can be tested without exposing individual cells to both treatments. This substantially increases the capabilities of the device for drug treatments, flux measurements, or local stimulation to address cell-cell communication and identify mechanisms of cell autonomy and intercellular coordination.

### Roots are able to sense asymmetric environments and adapt root hair development

To reveal how roots perceive and respond to environmental heterogeneity, we used the dfRootChip to provide an asymmetric micro-environment. Our data on fluorescein uptake demonstrated that external gradients of small molecules can be reflected by gradients throughout the root. In our dual flow perfusion system, such internal gradients are stabilised likely by active depletion of freely diffusing, apoplastic fluorescein molecules from the root side facing the flow of unstained solution.

We further tested whether asymmetric stimulation with stress elicitors results in asymmetric signalling. Our measurements on roots expressing R-GECO demonstrated that, as expected, calcium responses were first initiated on the side of stimulation. We observed, however, that the propagation of calcium signals differed substantially between responses to the biotic elicitor flg22 and the abiotic elicitor NaCl. While the response to salt stress propagated almost without attenuation across the root within less than 10 seconds (14.1 μm s^−1^), the elicited calcium response to flg22 treatment decayed beyond the vasculature and also propagated at an approximately 20 times slower rate (0.62 μm s^−1^). Recent studies used local application of salt solutions at the root tip and reported shootward long-distance propagation of calcium signals at approximately 400 μm s^−1^ (Choi et al., 2014; Evans et al., 2016). It was suggested that electrical signals may act as a fast mode of propagation along cells and the release of extracellular reactive oxygen species helps crossing cell-cell boundaries, albeit at likely a slower rate (Gilroy et al., 2014). The rate-limiting step would therefore be the number of cell-cell boundaries that are crossed over a distance. In line with this model, length:width ratios of fully elongated epidermis (13:1), cortex (10:1) and endodermis (18:1) cells (Hauser et al., 1995) could explain the slower rate of transverse versus longitudinal signal propagation. It remains, however, unclear why the calcium signals in response to flg22 only reached the vasculature and not the untreated side, while salt stress seemed to be perceived on both sides with little delay. A likely explanation for the localised calcium response to flg22 is the involvement of plasma membrane-based receptors and downstream signalling pathways, while NaCl may elicit calcium influx more directly due to mild osmotic effects. It can also be speculated that this behaviour reflects a fundamental difference between the two types of stresses, where biotic threats usually occur locally and activate local defence mechanisms, while abiotic stresses such as drought or salt affect the entire plant and require responses in distal cells to ensure survival of the organism. On the other hand, since responses to biotic stress often come at a significant cost to the plant, it may be advantageous to restrict responses of innate immunity to infected cells and tissues in their close vicinity.

When it comes to developmental adaptation of the root system architecture on the basis of nutrient availability the situation is more complicated. In soil environments with patchy distribution of barely diffusing nutrients, e.g. phosphate, evading depleted zones through growth requires some form of coordination as some cells potentially have to invest more resources than what can immediately be absorbed from the environment. Enhanced root hair growth is considered to be a typical adaptive response to low phosphate availability (Bates and Lynch, 1996; Karthikeyan et al., 2014; Ma et al., 2001; Song et al., 2016). The early finding that the transfer of seedlings between media with different phosphate content always triggered a phosphate concentration-specific response independent of the phosphate content in the medium before transfer was taken as evidence that root hair length is locally controlled (Bates and Lynch, 1996) and was later interpreted as a cell-autonomous response (Karthikeyan et al., 2014). However, so far no experiments had been performed where trichoblasts on the same root were exposed to different external phosphate concentrations at the same time. We therefore used root hair growth as a developmental readout under asymmetric phosphate supply. Instead of enhanced growth of root hairs, we observed, however, repressed root hair growth under phosphate deficient conditions. Although this was unexpected at first, it is in line with a study showing that hair growth stimulation under low phosphate depends on sucrose in the medium (Jain et al., 2007), which was part of the growth conditions in the aforementioned studies on root hair length regulation but absent in our media. We decided to not add sucrose to our media as we aimed to obtain root hair responses under conditions that approximate natural habitats where sucrose is also mostly absent. The apparent discrepancy between our results and previous work can also be explained by the influence of gelling agents on root hair growth, which was shown to have drastic effects with respect to phosphate-dependent hair growth regulation (Jain et al., 2009) and other root traits (Gruber et al., 2013). While previous work has usually employed agar-containing media to study root hair length under phosphate deficiency, our perfusion system involves exclusively synthetic hydroponic media. An additional concern, regarding root hair measurements on vertical plates is the inherent asymmetry in the experimental growth conditions, with one side of the root being in direct contact with the substrate, while the other side is exposed to air. Recent work demonstrated that hair growth is stimulated on the air side and repressed on the medium side (Bao et al., 2014), which means that hair length measurements in many previous phosphate deficiency experiments may have mostly evaluated trichoblasts that were not in direct contact with the medium.

Our experiments, using hydroponic perfusion in the dfRootChip demonstrated that root hair growth under phosphate-deficient conditions can be repressed, independent of the overall phosphate supply. This local response is evidence for a cell- or cell-file-autonomous negative regulation of root hair growth. Yet, hair growth on the phosphate-sufficient side was stimulated by approximately 32% when the opposite side of the root was exposed to phosphate-deficient conditions. This response of distal cells indicates the existence of intercellular communication in order to positively regulate root hair growth. Ethylene signalling could present a potential mechanism for this intercellular regulation, since recent studies showed an involvement in root hair growth control under phosphate deficient conditions (Nagarajan et al., 2011; Song et al., 2016; Song and Liu, 2015). Interestingly, a very similar response was observed under asymmetric water-stress conditions; high osmolarity of the perfusion medium containing 20% PEG (w/v) resulted in hair repression and an approximately 21% stimulation of hair growth on the opposite side of the root. We therefore conclude that root hair development can be regulated on both, the cellular and the organ level. Future studies will unveil the mechanisms and regulatory networks that allow plants to adapt their root morphology.

### Conclusions and perspectives

Environmental complexity will remain a major challenge for studies on plant signalling and development. Yet, we cannot ignore this characteristic feature of natural plant habitats when aiming to understand plant development of model and crop plants alike (Rellán-Álvarez et al., 2016). Advanced technologies for plant cultivation, imaging and quantification will enable us to better mimic field conditions in the laboratory and thereby provide the basis for new discoveries (Ehrhardt and Frommer, 2012). In recent years, the adaptation of microfluidics for organismal studies has demonstrated great potential to create dynamic micro-environments and aid new experimental approaches for a number of model organisms (Stanley et al., 2015). It will be interesting to systematically test combinations of environmental conditions in the dfRootChip and reveal further developmental trade-offs and decision-making. Particularly the visualisation of proteins and small molecules using genetically encoded nanosensors is benefitting from the defined conditions provided by Lab-on-a-Chip devices (Jones et al., 2013; Uslu and Grossmann, 2016). In combination, nanosensors and microfluidics offer countless possibilities for studies on plant-environment interactions including tracing nutrient absorption, understanding environmental sensing and stress signalling or investigating intercellular and inter-organismal communication. With environmental complexity taken into account, these studies will yield novel molecular mechanisms and genetic traits that will benefit crop breeding and field research.

## MATERIALS AND METHODS

### Plant lines and growth conditions

*Arabidopsis thaliana* seeds (Col-0 and R-GECO) were prepared for imaging as described previously (Grossmann et al., 2012) with some modifications detailed herein. Briefly, the seeds were surface sterilised using 5% sodium hypochlorite solution (Sigma Aldrich, Germany) by vortexing the seeds for 3-5 mins and rinsing three times with sterile double distilled water. The seeds were stratified for 3 days at 4 °C and germinated on 10 μl pipette tips (cut to ca. 5 mm in length) filled with half strength Hoagland’s medium (1/2x HM) solidified with 0.7% (w/v) plant agar (Duchefa Biochemie, Germany) (Supplementary Table S1). In all experiments the plants were grown continuously under long day conditions (16 h light, 22 °C, 100 μΕ m^−2^ s^−1^, 50-70% relative humidity) except for those involving poly(ethylene glycol) (PEG) treatments, which were grown under short day conditions (9 h light, 24 °C, 180 μΕ m^−2^ s^−1^, 53% relative humidity).

### Microfluidic device design and mould fabrication

The device design was constructed in AutoCAD Mechanical 2011 (Autodesk) and used to create a mylar film photolithography mask (Micro Lithography Services Ltd., UK). The master mold was manufactured using photolithography (Duffy et al., 1998). Briefly, a 100 mm silicon wafer (Silicon Materials, Germany) was spin-coated with SU-8 3050 photoresist (MicroChem, USA) by dispensing ca. 4 ml of photoresist onto the wafer while spinning at 100 rpm for 5 s with acceleration of 10 rpm/s, then spinning at 500 rpm for 10 s with acceleration of 5 rpm/s and finally at 500 rpm for 10 s with acceleration of 5 rpm/s. The resist was baked at 95°C for 20 minutes for hardening and then exposed in an MA6 ultraviolet (UV) mask aligner (Suss Microtec, Germany) using the mylar film photolithography mask, with an exposure energy dose of 500 mJ/cm^2^ (λ = 365 nm). The resist was again baked at 95°C for 12 minutes. Finally, the SU-8 resist was developed using mr-Dev 600 developer solution (Microresist Technologies, Germany) and characterised using a Dektak XT stylus profilometer (Bruker, USA). The height of the SU-8 structures was found to be on average 115 ± 10 μm in height. The master moulds were then silanised under vacuum for 2 hours with 50 mL chlorotrimethylsilane (Fluka, Germany) per master.

### Device fabrication

PDMS was prepared using a 10:1 ratio of base to curing agent (Sylgard 184, Dow Corning, USA). The base and the curing agent were mixed together thoroughly, degassed for 1 hour under vacuum and poured on top of the master mold. Cover slips (45 × 50 mm, #1, Menzel GmbH, Germany) were spin coated (Laurell Technologies, USA) with approx. 0.02 mm PDMS. The PDMS was then cured overnight at 70 °C, removed from the master mold and diced to size. Precision cutters (Syneo, USA) were used to punch holes to form the root inlets (1.65 mm cutting edge diameter), solution in- and outlets (1.02 mm cutting edge diameter) and reservoirs (4.75 mm cutting edge diameter). The PDMS top layer, containing the microchannels, as well as the PDMS-coated cover slip, were washed and dried as detailed elsewhere (Stanley et al., 2014). Bonding of the PDMS top layer was sealed to the bottom layer(s) of the device using a Diener ZEPTO plasma cleaner (Diener electronic, Germany; conditions: power 50%; 1 min.)

### On-chip plant cultivation, media perfusion and treatments

Microfluidic devices were prepared as described previously (Grossmann et al., 2012; 2011) with minor modifications. Briefly, the devices were sterilised under ultraviolet light for 20 minutes and the microchannels were filled with 1/2x HM by manually pipetting medium through the root inlet. The inlets and reservoirs were then topped up with 1/2x HM to ensure that they were completely filled. The Arabidopsis seedlings were visually inspected using a stereoscope 4-5 days after germination to select for roots being at approximately the same growth stage. The pipette tips containing the Arabidopsis plants were then inserted into the root inlets under sterile conditions. Each device was then placed into a round plastic petri dish, ca. 15 ml of 1/2x HM introduced to provide a humid environment and each petri dish sealed with parafilm. The petri dishes containing the microfluidic devices and plants were then transferred into the growth chamber, tilted at an angle of ca. 20° to encourage root growth in the direction of the microchannels and left until the roots had grown into the main observation channels. The device was then transferred to a homemade rectangular chip carrier, fixed with an adhesive tape and subsequently mounted on the microscope stage. To maintain a humid environment, the dfRootChip was surrounded by moist tissues and covered with a transparent plastic lid.

### Poly(ethylene glycol) (PEG) experiments

A 20% (w/v) PEG solution was prepared as follows. 40 g of PEG 8000 (Sigma-Aldrich, USA) was dissolved in 100 ml of autoclaved full strength Hoagland’s Medium (Supplementary Table S1) and made up to a final volume of 200 ml with double distilled water. The pH was adjusted to 5.7. PEG solutions were always prepared freshly and sterile filtered (0.22 μm filter) before use. Symmetric and asymmetric treatments were performed using two Aladdin-220 syringe pumps (World Precision Instruments, UK) to pump media into the microchannels. Specifically, two 60 ml BD Plastipak sterile syringes containing the test solutions were held in the syringe pump holders and each connected to a 7-port manifold (P-150, Ercatech AG, Switzerland) using a flangeless fitting for 1/16” tubing, luer adapter (Ercatech AG, Switzerland) and polytetrafluoroethylene (PTFE) tubing with inner and outer diameters of 0.25 mm and 1/16” respectively (Upchurch Scientific, Germany). Each manifold was used to split the input flow into five separate flows, therefore serving five inlet channels (either inlets 1A-5A or 1B-5B, Supplemental Fig. 1). Tubing from the manifold was directly inserted into the device inlets. Before each experiment, the tubing was thoroughly rinsed with 70% ethanol or autoclaved. Connection of the tubing to the device inlets was always conducted in a sterile hood and fluid was pumped through the tubing when connecting them to the device inlets to prevent the introduction of air bubbles. The flow rate per channel was set to 8 μl/min. Treatments were conducted immediately after removing the microfluidic devices from the plant growth chamber.

### Phosphate experiments

For perfusion experiments under phosphate rich (2.5 mM KH_2_PO_4_) or phosphate deficient (0.01 mM KH_2_PO_4_ supplemented with 2.49 mM KCl) conditions, liquid media were prepared as described elsewhere (Chandrika et al., 2013) without the addition of sucrose (**Supplementary Table S1**). For experiments performed on agar plates, the same media were supplemented with 0.7% plant agar (Duchefa, Germany). Treatments were performed as detailed for PEG experiments with the following modifications. The device was connected to a six-channel syringe pump (NE-1600, New Era Pump Systems, USA) via Tygon flexible plastic tubing (ID 0.02 inch; OD 0.06 inch; Saint-Gobain Performance Plastics, France) and stainless steel connectors (0.025” OD, 0.013” ID, 0.5” long; New England Small Tube, USA). The flow rate was set to 5 μl/min per inlet channel. Initially, roots were perfused with phosphate rich medium on both sides until the root tips had entered the observation channels. The phosphate rich medium was then i) continued or ii) switched on either one or both sides of the roots to perform an asymmetric or symmetric perfusion with the deficient phosphate medium respectively.

### Rapid asymmetric treatments with calcium elicitors

A pressurised vial (10 ml volume) containing the 1/2x HM was connected to one of the device inlets via Tygon tubing, stainless steel connectors and a luer-lock stopcock valve (Vygon, Germany) set to an “open” configuration. The control solution was introduced into the microchannels using a volumetric flow rate of 20 μl/min until the solution exited the outlet and second solution inlet. A second tubing prefilled with treatment solution (1/2x HM containing either 100 mM NaCl or 1 μΜ flg22) was connected to a second pressurised vial via a stopcock valve in a “closed” configuration. This treatment tubing is then connected to the second solution inlet of the dfRootChip. The actively flowing control solution prevents the treatment solution from entering into the device before the stopcock valve is opened. Imaging is started and the treatment solution valve is opened at a given time point to initiate immediate treatment. The control valve is kept open to achieve asymmetric treatment. If symmetric treatment with the treatment solution is required, the control valve is closed and treatment valve opened simultaneously.

### Treatments with *Pseudomonas fluorescens*

The bacterial strain *Pseudomonas fluorescens* WCS365 carrying a plasmid for expression of green fluorescent protein (GFP) (Haney et al., 2015) was maintained on lysogeny broth (LB) medium supplemented with 50 μg/ml kanamycin and grown aerobically at 28 °C. For experiments, WCS365 cells were inoculated in 20 ml LB supplemented with kanamycin (50 μg/ml), pelleted during the exponential growth phase (OD_600_ = 0.1-0.2) and resuspended in 20 ml 1/2x HM. The *P. fluorescens* cell suspension was directly used for treatments of RGECO roots in the dfRootChip. Before treating the roots with *P. fluorescens*, they were first perfused with 1/2x HM until the root tips had entered the observation channels. The perfusion set-up used in this experiment was the same as described for the phosphate experiment. The flow rate was set to 5 μl/min per inlet channel for the duration of the experiment.

### Imaging and image analysis

Bright field and fluorescence images as well as time series of growing plant roots were taken on a Nikon Ti-U inverted microscope, equipped with a x10 air immersion N.A. 0.30 objective (Nikon, Switzerland), a Prior ProScan III motorised stage (Prior Scientific, UK) and a CoolSNAP HQ2 camera (Photometrics, Germany) or a Nikon Ti-E, equipped with 20x air immersion N.A. 0.7 and 60x water immersion N.A. 1.2 objectives (Nikon), a motorized XYZ stage (Applied Scientific Instrumentation, USA), a filter wheel (Cairn Optospin, UK), a laser launch (Omicron, Light Hub, Germany) housing 5 laser lines (440, 488, 515, 561, 638 nm), and an EMCCD camera (Photometrics, Evolve Delta, USA). NIS-Elements Advanced Research imaging software (Nikon, Switzerland) with autofocus or Micro-Manager 1.4 were used to synchronise long-term, multi-position, time-lapse imaging experiments at an ambient temperature of 24°C. The images were analysed using Fiji (Schindelin et al., 2012). Root hair length was measured using the freehand line tool in Fiji and an Intuos pen tablet (Wacom, USA). Statistical significance between data sets was calculated using a Student’s *t*-test. Root hair growth within ca. 1500 μm (phosphate experiments) or 1200 μm (PEG experiments) of the growing root tip was excluded during the analysis to discount newly elongating or emerging root hairs. Box blot graphs were generated in R with the help of the BoxPlotR web tool (http://shiny.chemgrid.org/boxplotr/). The authors are grateful to the Tyers and Rappsilber labs for providing this helpful resource.

## SUPPLEMENTARY INFORMATION

### Supplementary Figures

**Supplementary Figure S1.**
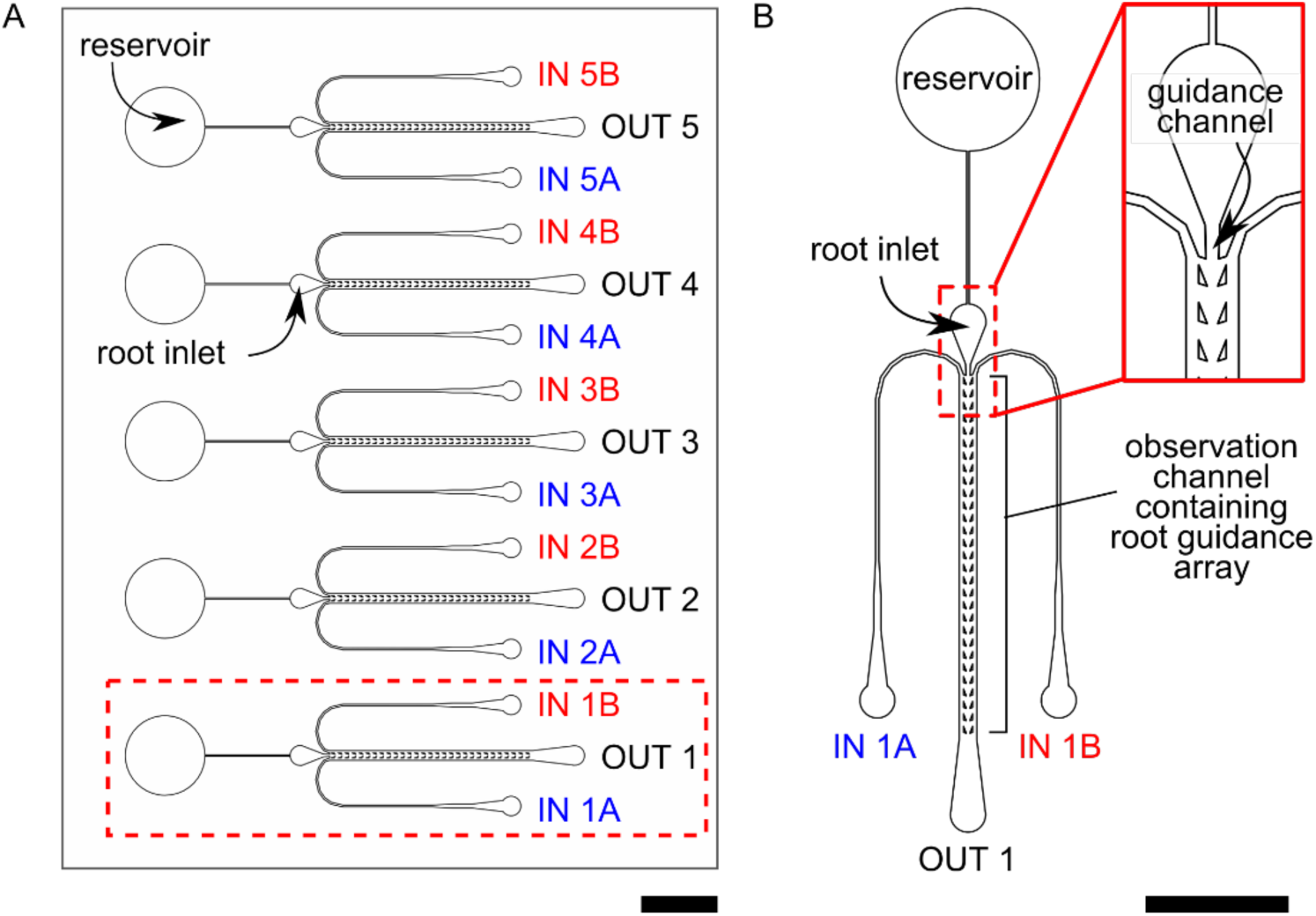
Design master for the dual-flow-RootChip (dfRootChip). (**A**) Overview of the design master for the dfRootChip, in which a plant chamber (highlighted by the dotted red box) can be replicated to enable several plant root experiments to be conducted in parallel. We exemplify this by combining five chambers in parallel on a single glass slide. The dfRootChip consists of ten inlet channels (two serving each plant chamber) and five outlets. A plant growth medium reservoir is connected to each root inlet (and consequently to the plant root observation channel) via a small channel. (**B**) An enlarged version of a single plant chamber, where the reservoir, root inlet containing a guidance channel (dotted red box), observation channel and root guidance array are highlighted. Scale bars represent 4 mm.

**Supplementary Figure S2.**
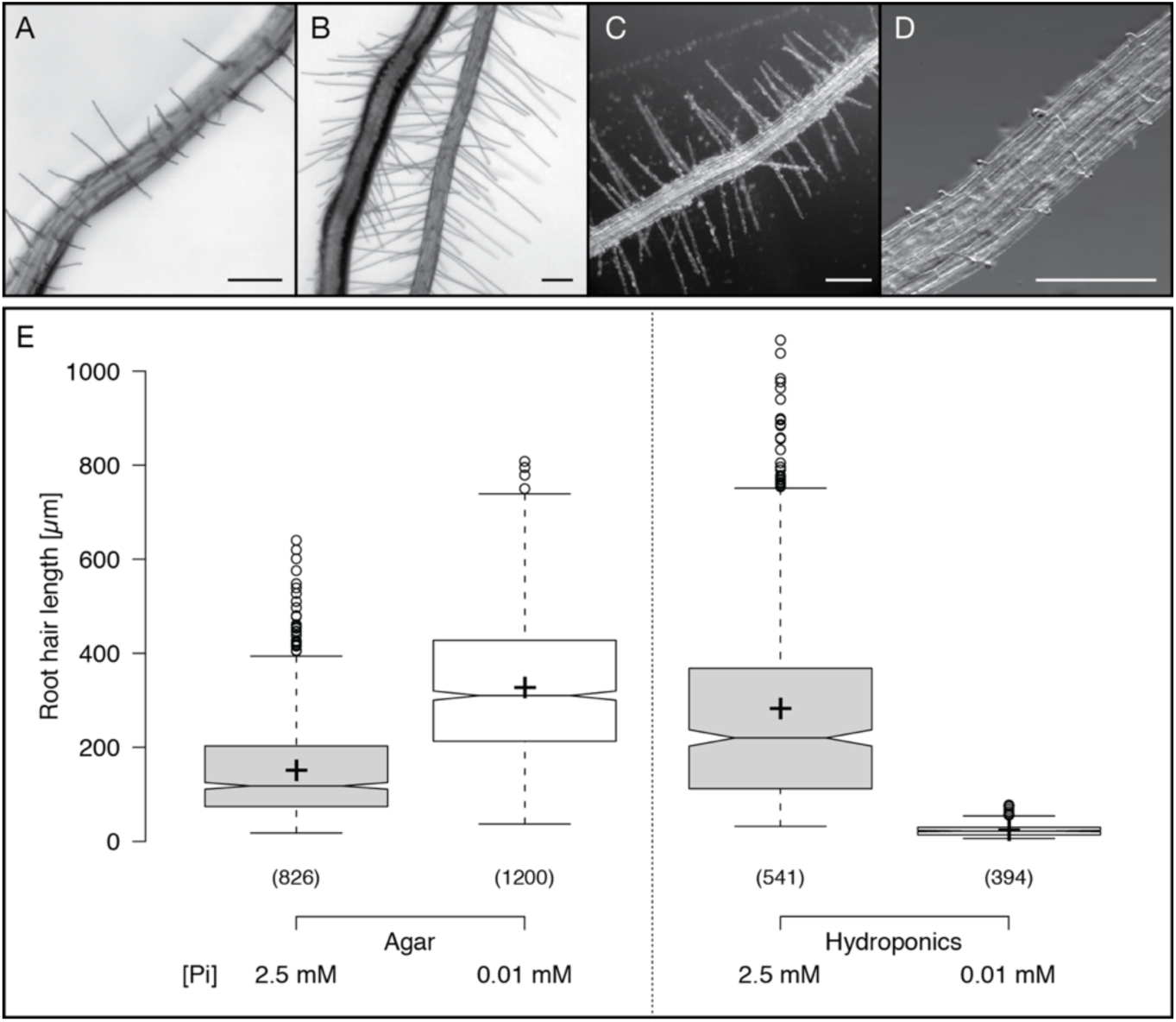
Root hair growth under high and low phosphate availability on agar plates and in hydroponic medium. (**A-D**) Representative images of primary roots of 10 days old Col-0 seedlings grown on solid medium containing (**A**) 2.5 mM KH_2_PO_4_, (**B**) 0.01 mM KH_2_PO_4_ and in hydroponic culture containing (**C**) 2.5 mM KH_2_PO_4_ and (**D**) 0.01 mM KH_2_PO_4_. All scale bars represent 100 μm. (**E**) Quantification of root hair length for the conditions given in (**A-D**). Centre lines depict the median values, crosses represent sample means; box limits indicate the 25th and 75th percentiles; whiskers extend 1.5 times the interquartile range from the 25th and 75th percentiles, outliers are represented by dots. Sample points (n) from at least 2 independent experiments (agar plates) and 3 independent experiments (hydroponics), respectively, are given in parentheses below each box.

### Supplementary Movie Legends

**Supplementary Movie 1.** An Arabidopsis root growing in the dfRootChip. The time interval between successive frames is 2 minutes. The playback rate is 20 fps. Scale bar, 200 μm.

**Supplementary Movie 2.** Laminar flow in the dfRootChip. Two co-flowing liquids (water), one of which was supplemented with food colouring to visualise flow separation. Bright field images were captured and false-coloured using the “Royal” lookup table. At time point 8.9 s, the flow pressure is changed from 25 mbar to zero, resulting in a breakdown of laminar flow. The playback rate is 20 fps (real-time playback). Scale bar, 200 μm.

**Supplementary Movie 3.** Normalised R-GECO fluorescence images (ΔF/F) of calcium dependent signal changes upon asymmetric stimulation with 1 μΜ flg22 in 1/2x HM. The treatment was introduced on the right side. The time interval between successive frames is 0.75 s. The playback rate is 60 fps. Scale bar, 100 μm.

**Supplementary Movie 4.** Normalised R-GECO fluorescence images (ΔF/F) of calcium dependent signal changes upon asymmetric stimulation with 100 mM NaCl in 1/2x HM. The treatment was introduced on the right side. The time interval between successive frames is 0.75 s. The playback rate is 60 fps. Scale bar, 100 μm

### Supplementary Materials and Methods

**Supplementary Methods S1 | Phosphate experiments conducted in containers**

For hydroponic phosphate experiments conducted in small containers, the phosphate rich and deficient media were prepared as described in Supplementary Table S1. Five-day-old Col-0 seedlings grown in pipette tips filled with 1/2x HM medium were inserted into holes punched into a thin PDMS slab, which was then mounted on a round float rack (VWR, America). This set-up was transferred to sterivent high containers (107 × 94 × 96 mm; Duchefa, Germany) filled with 500 ml of either phosphate rich or deficient medium under sterile conditions and the seedlings grown under long day conditions (16 h light, 22 °C, 100 μΕ m^−2^ s^−1^, 50-70% relative humidity) for 5-7 days. To acquire images, roots were cut, and gently placed on flat glass slides (76 × 26 mm; Labsolute, Germany) and a cover glass slide (24 × 50 mm; Roth, Germany) laid on top. Images were acquired using a SMZ18 Nikon stereoscope (Nikon, Japan) equipped with a 2x objective and Orca Flash 4.0 camera (Hamamatsu, Japan).

**Supplementary Table S1.** Media used in this study.

| **Medium** | **Details** |
| --- | --- |
| 1/2x Hoagland's medium (1/2x HM) | 0.8 g/L Hoagland's no. 2 basal salt mixture, 0.5 g/L MES hydrate. Adjusted to pH 5.7 with KOH. Agar plates made with 1/2x HM were solidified with 0.7% w/v agar. |
| Medium containing Poly(ethylene glycol) | 1/2x HM containing 20% w/v PEG 8000. Adjusted to pH 5.7 with KOH. |
| Media containing Ca<sup>2+</sup> elicitors | 1/2x HM containing either 100 mM NaCl or 1 μM flg22 |
| Phosphate rich medium<br>(Chandrika et. al. 2013) | 2.5 mM K<sub>2</sub>H<sub>22</sub>PO<sub>4</sub>, 2 mM MgSO<sub>4</sub>·7H<sub>2</sub>O, 5 mM KNO<sub>3</sub>, 2 mM Ca(NO<sub>3</sub>)<sub>2</sub>·4H<sub>2</sub>O, 0.04 mM Na-Fe-EDTA, 70 μM H<sub>3</sub>BO<sub>3</sub>, 0.5 μM CuSO<sub>4</sub>·5H<sub>2</sub>O, 1 μM ZnSO<sub>4</sub>·7H<sub>2</sub>O, 14 μM MnCl<sub>2</sub>·2H<sub>2</sub>O, 0.2 μM Na<sub>2</sub>MoO<sub>4</sub>, 0.01 μM CoCl<sub>2</sub>·6H<sub>2</sub>O, 1 g/L MES hydrate. Adjusted to pH 5.5 with KOH. Agar plates made with this medium were solidified with 0.7% w/v agar. |
| Phosphate deficient medium<br>(Chandrika et. al. 2013) | 0.01 mM KH<sub>2</sub>PO<sub>4</sub>, 2.49 mM KCl, 2 mM MgSO<sub>4</sub>·7H<sub>2</sub>O, 5 mM KNO<sub>3</sub>, 2 mM Ca(NO<sub>3</sub>)<sub>2</sub>·4H<sub>2</sub>O, 0.04 mM Na-Fe-EDTA, 70 μM H<sub>3</sub>BO<sub>3</sub>, 0.5 μM CuSO<sub>4</sub>·5H<sub>2</sub>O, 1 μM ZnSO<sub>4</sub>·7H<sub>2</sub>O, 14 μM MnCl<sub>2</sub>·2H<sub>2</sub>O, 0.2 μM Na<sub>2</sub>MoO<sub>4</sub>, 0.01 μM CoCl<sub>2</sub>·6H<sub>2</sub>O, 1 g/L MES hydrate. Adjusted to pH 5.5 with KOH. Agar plates made with this medium were solidified with 0.7% w/v agar. |
| Lysogeny broth (LB) medium | 1% w/v tryptone, 0.5% w/v yeast extract, 1% w/v NaCl |

## ACKNOWLEDGEMENTS

We thank Cara Haney (Vancouver) and Ikram Blilou (Wageningen) for sharing fluorescent *Pseudomonas* strains. We are grateful to Apolonio Huerta for his assistance in device fabrication, to Karin Schumacher and Rubén Rellan Alvarez for critical reading of the manuscript and to the members of the Grossmann lab for inspiring discussions. We also thank Andrew deMello and members of the deMello Lab for discussions and finally Justina Rutkauskaite for the beautiful photograph illustrated in Figure 1A. We acknowledge financial support by the Eidgenössische Technische Hochschule Zürich (CES), a Faculty for the Future Fellowship by the Schlumberger Foundation to JS and the Heidelberg Excellence Cluster CellNetworks to GG.

## AUTHOR CONTRIBUTIONS

The study was conceived by GG, the device was designed by CES and GG, with help from DS. CES and JS performed fluorescein, growth and root hair measurements. Experiments involving fluorescent bacteria were conducted by JS. JS and RB performed calcium measurements. CES, JS, RB and GG analysed the data. CES and GG wrote the manuscript with input from JS and RB. All authors read and approved the final version of the manuscript.

